# X-ray crystallographic fragment screening reveals novel and conformationally dynamic ligand-binding sites in *Mycobacterium tuberculosis* FtsZ

**DOI:** 10.64898/2026.09.01.748605

**Authors:** Kaylen R. Meeks, Edwin O. Lazo, Dale F. Kreitler

## Abstract

Tuberculosis is a leading cause of death globally due to an infectious agent. There is ongoing need for novel mechanisms to inhibit *M. tuberculosis* (Mtb) growth and infection to improve patient outcomes. FtsZ, a GTPase that assembles into protofilaments at the division site of a replicating cell to produce two individual cells, is an attractive target as an essential protein in bacterial cell division. Here we describe a crystallographic fragment screening campaign of MtbFtsZ. 1,070 crystals were soaked with fragments and 714 datasets were used for downstream PanDDA analysis. 149 datasets exhibited PanDDA-generated event map density to support modeling of fragment binding. 15 novel sites are described. Both the ON and the OFF conformations of FtsZ are found in the asymmetric unit. Asymmetric binding of fragments to each chain in the model is observed. These crystallographic fragment screening results additionally provide opportunities for fragment growing and merging to develop FtsZ binders into drug-like molecules or conformation specific chemical probes.

## Introduction

Tuberculosis (TB) is one of the leading causes of death worldwide due to an infectious agent, causing 1.23 million deaths in 2024, according to the World Health Organization^1,2^. TB is contracted after infection by the pathogenic bacterium *Mycobacterium tuberculosis*^3^. Strains of *M. tuberculosis* that do not respond to first-line drugs, commonly referred to as multidrug-resistant (MDR-TB), have introduced challenges to TB treatment. MDR-TB requires more extensive and expensive care than conventional TB with current treatment strategies^3,4^. Modern antibiotics, such as pretomanid and bedaquiline, have shown promise in treating MDR-TB; although, single nucleotide polymorphisms have conferred resistance to drugs, including bedaquiline^3,5–7^. Therefore novel therapeutics are urgently needed to revolutionize TB care^5^.

Filamentous temperature-sensitive mutant Z (FtsZ) is critical for *M. tuberculosis* cell division^8,9^. FtsZ polymerizes in a GTP-dependent manner to form the Z-ring, which assembles at the division site and provides a scaffold for the recruitment of additional components of the divisome^8^. In mycobacteria, FtsZ interacts with several cell-division proteins including SepF, FtsW, and CrgA^10,11^. SepF contributes to Z-ring assembly and localization at the membrane, whereas interactions with FtsW and CrgA provide connections between the Z-ring and downstream components involved in septal peptidoglycan synthesis. FtsZ has gained broad interest as an antibacterial drug target due to its essential role in bacterial cell division and its interaction with multiple components of the divisome.

MtbFtsZ crystallizes as an asymmetric dimer containing two distinct conformational states (Fig. 1A)^12^. In the GTPγS-bound protomer (chain A), the T3 loop is ordered upon γ-phosphate binding, which stabilizes switch helix 2 (sH2) in its α-helical conformation, as the β-strand conformation (sβ2) would clash with the ordered T3 loop. Together, the ordered T3 loop and α-helical sH2 define the GTP-associated “ON” configuration. In contrast, chain B adopts an “OFF” configuration in which the nucleotide binding site is occluded, sH2 adopts the β-strand conformation (sβ2), and the T3 loop is disordered. Superposition of the two protomers highlights these conformational differences, including rearrangements of sH2/sβ2 and the central helix H8 (Fig. 1B). Together with the central helix (H8), these structural elements couple nucleotide binding to conformational changes associated with FtsZ function. The C-terminal T7 loop contributes to the longitudinal protofilament assembly by extending into the nucleotide-binding pocket of the adjacent subunit, where it participates in GTP hydrolysis^12–15^.

**Figure 1.**
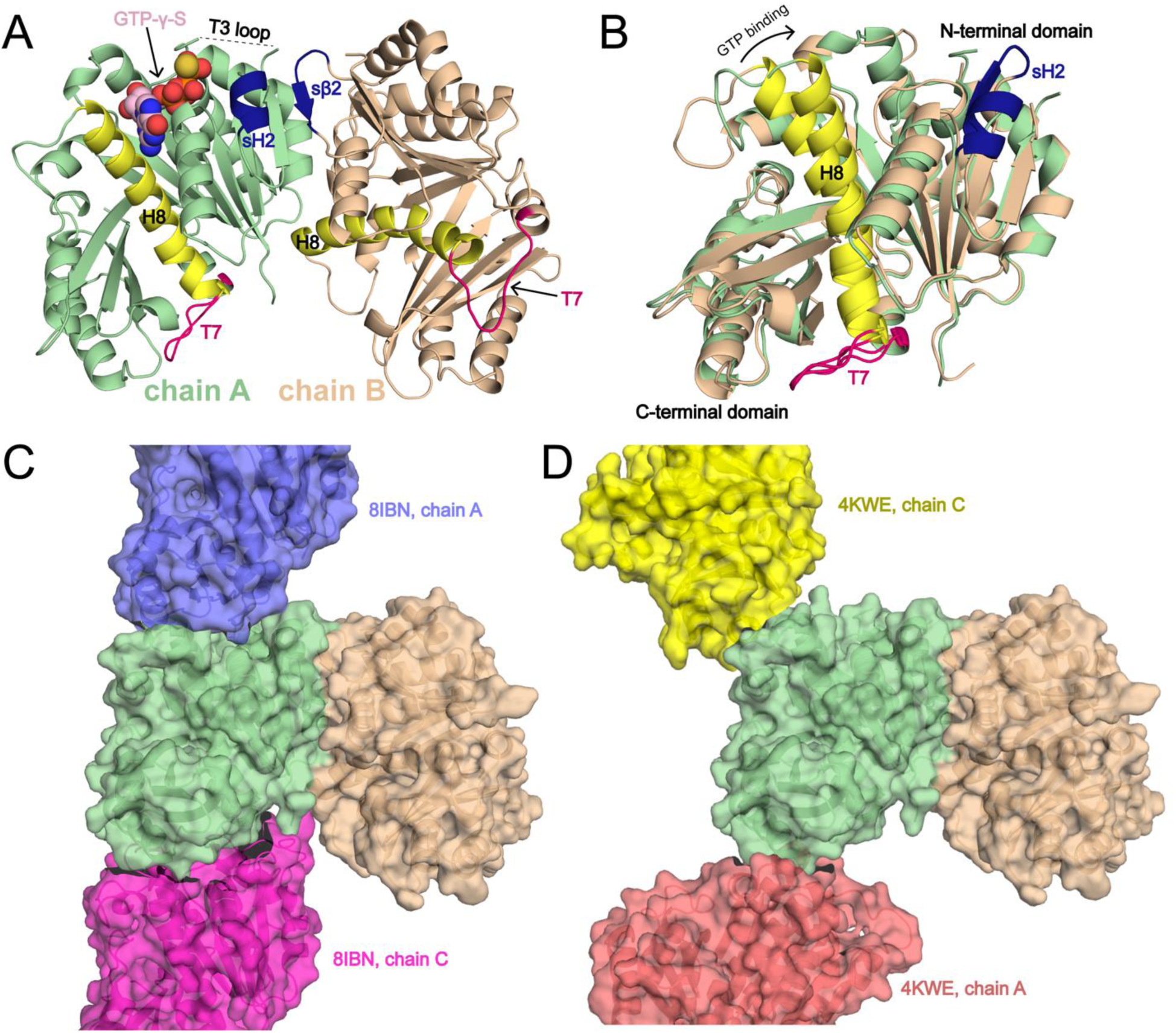
FtsZ secondary structure and polymerization. **A**) The ground state (pdb_000015aa) FtsZ model is shown in cartoon representation, with chain A in pale green (ON protomer) and chain B (OFF protomer) in wheat. Key structural elements associated with the conformational state and protofilament assembly of FtsZ are highlighted: helix 8 (H8, yellow), T7 loop (T7, dark pink), switch helix 2 (sH2, dark blue), and T3 loop. The T3 loop is disordered in the ground-state model but ordered in the GTPγ-S-bound structure pdb_00001rlu. **B**) Superposition of Chain A and chain B of the ground-state model aligned using H8. **C**) Cryo-EM structure of KpFtsZ filament (pdb_00008ibn) overlayed on MtbFtsZ ground state model. The ground state model is shown in pale green and wheat. The KpFtsZ filament chains are shown in purple and pink. **D**) Crystal structure of MtbFtsZ curved filament (pdb_00004kwe, red and yellow) overlayed on MtbFtsZ ground state model.

Structural studies of FtsZ from multiple species have revealed straight and curved protofilament conformations associated with its dynamic role in Z-ring assembly and constriction^15–17^. A straight protofilament conformation is represented by the *Klebsiella pneumoniae* FtsZ (KpFtsZ) structure (pdb_00008ibn; Fig. 1C)^16^, whereas a GDP-bound MtbFtsZ structure captures a curved protofilament conformation (pdb_00004kwe; Fig. 1D)^15^.Together, these structures illustrate the conformational plasticity of FtsZ and provide a framework for evaluating whether ligand-binding sites identified in this study overlap structural elements involved in protofilament assembly and conformational remodeling.

Several strategies have been explored to inhibit FtsZ, including disruption of protofilament assembly, modulation of filament dynamics, and competition with nucleotide binding. White *et al.* screened a series of 2-alkoxycarbonylaminopyridines originally developed as potential tubulin (the eukaryotic homolog of FtsZ) inhibitors and identified SRI-3072 (Pubchem: 493469) as a selective inhibitor of MtbFtsZ filament polymerization (ID_50_ = 52 ± 12 μM) relative to tubulin^9,18^. Subsequent attempts to obtain structural evidence for binding of the related compound SRI-7614 (Pubchem: 493470) to MtbFtsZ were unsuccessful^19^. In contrast, Alnami *et al.* demonstrated crystallographically that 4-hydroxycoumarin binds two cryptic pockets in MtbFtsZ^13^. Although these structures provide a mechanistic basis for inhibition, quantitative binding affinity and direct experimental validation linking occupancy of these cryptic pockets to MtbFtsZ inhibition have not been reported^13^.

Allosteric inhibitors of FtsZ have also been characterized in other bacterial species, particularly *Staphylococcus aureus*^20–24^. TXH9179 (Pubchem: 167312464) binds a hydrophobic pocket in SaFtsZ adjacent to the T7 loop and located between the core helix (H8 in MtbFtsZ) and the C-terminal β-sheet (Fig. S4)^23,24^. Structural comparison with MtbFtsZ shows that its C-terminal β-sheet is positioned ∼10 Å closer to H8, substantially reducing the volume of the corresponding pocket (Fig. S4A). A similarly restricted pocket is observed in *Escherichia coli* FtsZ (EcFtsZ) structures^25^. Nevertheless, BOFP (Pubchem: 145945984), a compound known to bind this pocket, has micromolar affinity for EcFtsZ in fluorescence anisotropy experiments^26^. These observations highlight the difficulty of predicting FtsZ ligandability from individual static structures and motivate direct experimental identification of ligand-binding sites in MtbFtsZ.

We used high-throughput x-ray crystallographic fragment screening to discover novel binding pockets of MtbFtsZ. Fragment screening uses low molecular weight (<250Da) compounds to sample a diverse chemical space. Fragments typically have weak affinity for a given target macromolecule, because their small size limits the number of favorable interactions between fragments and the target. However, weak and oftentimes promiscuous binding is advantageous for identifying regions of a protein that are capable of ligand binding^27–29^. X-ray crystallography is well suited for fragment screening because it directly identifies binding sites and ligand poses, providing structural information for subsequent fragment growing, merging, or linking^27,30^. Eight FDA-approved drugs have been discovered through fragment-based approaches, with more in clinical trials^27,28^. Advances in automated data collection at modern synchrotron beamlines, such as the Highly Automated Macromolecular Crystallography (AMX) beamline at the National Synchrotron Light Source II (NSLS-II), have made x-ray crystallographic fragment screening a viable primary screening method^31,32^.

Here, we report a high-throughput x-ray crystallographic fragment screen of MtbFtsZ that identifies 20 ligand-binding sites distributed across the protein, including 15 sites not previously described in MtbFtsZ. The presence of conformationally distinct ON and OFF protomers within the asymmetric unit further enabled direct comparison of fragment binding between two structural states, revealing pronounced state-dependent differences in pocket ligandability. Together, these structures substantially expand the experimentally defined ligand-binding landscape of MtbFtsZ and provide starting points for chemical interrogation of nucleotide-binding, conformationally dynamic, and protofilament-associated regions of the protein.

## Results

### MtbFtsZ crystals are suitable for high-throughput fragment screening

MtbFtsZ was screened for crystallization conditions using Index Screen (Hampton Research). Initial crystals obtained in condition F6 condition (0.2 M Ammonium sulfate, 0.1 M Bis-Tris pH 5.5, 25% (w/v) PEG 3,350) were further optimized using microseeding. Under the optimized conditions, MtbFtsZ crystallized primarily in space group *P*6_5_ with two protomers in the asymmetric unit. This crystal form is consistent with previously reported MtbFtsZ structures such as pdb_00001rlu and related entries^32^.

The two protomers adopt distinct conformational states corresponding to the GTP-associated MtbFtsZ ON and OFF states (Fig. 1A-B). This crystal form therefore provided an opportunity to simultaneously evaluate fragment binding to two structurally distinct states of the protein within the same crystal lattice.

Prior to fragment screening, the tolerance of MtbFtsZ crystals to DMSO and extended soaking was evaluated using 32 crystals with soak times ranging from 80 to 125 min. Twenty-eight datasets had an average high-resolution limit of 2.3 ± 0.4 Å, with no apparent relationship between soaking time and diffraction resolution. The tolerance of the crystals to DMSO soaking while maintaining diffraction at approximately 2-3 Å resolution supported their use for high-throughput x-ray crystallographic fragment screening.

### Crystallographic fragment screening identifies 149 hits distributed across 20 binding sites

Crystallographic fragment screening was performed using the DSi-Poised (Enamine, Kyiv, UA) and Fragment Diversity Set III (Life Chemicals, Niagara-on-the-Lake, CA) libraries. A total of 1,070 fragment-soaked crystals were harvested and 843 diffraction datasets were obtained. Following exclusion of datasets because of space-group heterogeneity, resolution, or poor processing, 714 datasets were included in PanDDA analysis. The datasets had an average high-resolution limit of 2.29 ± 0.36 Å (Fig. S1).

Manual inspection of PanDDA event maps identified 149 datasets containing electron density supporting placement of one or more fragments, corresponding to an overall crystallographic hit rate of 21%. Forty hit datasets (27% of hits) contained multiple poses of the same fragment within a single structure at different binding sites. The average resolution of the hit datasets was 2.08 ± 0.22 Å. Ensemble models representing both the ground and fragment-bound states were generated for refinement, resulting in a mean *R*_free_ of 0.22 ± 0.01 (Fig. S1).

Spatial grouping of the fragments identified 20 distinct binding sites distributed across the MtbFtsZ structure (Fig. 2 and S3). Ten sites contained at least three overlapping fragments and were classified as fragment clusters. Five of the observed sites correspond to previously described ligand-binding regions, including the ON and OFF nucleotide-binding pockets (sites 6 and 7), cryptic pocket 1 (site 3), and cryptic pocket 2 in the two protomers (sites 1 and 4). The remaining 15 sites have not previously been described as ligand-binding sites in MtbFtsZ. The locations, conformational states, fragment occupancy, and structural context of all 20 sites are summarized in Table 1.

**Figure 2.**
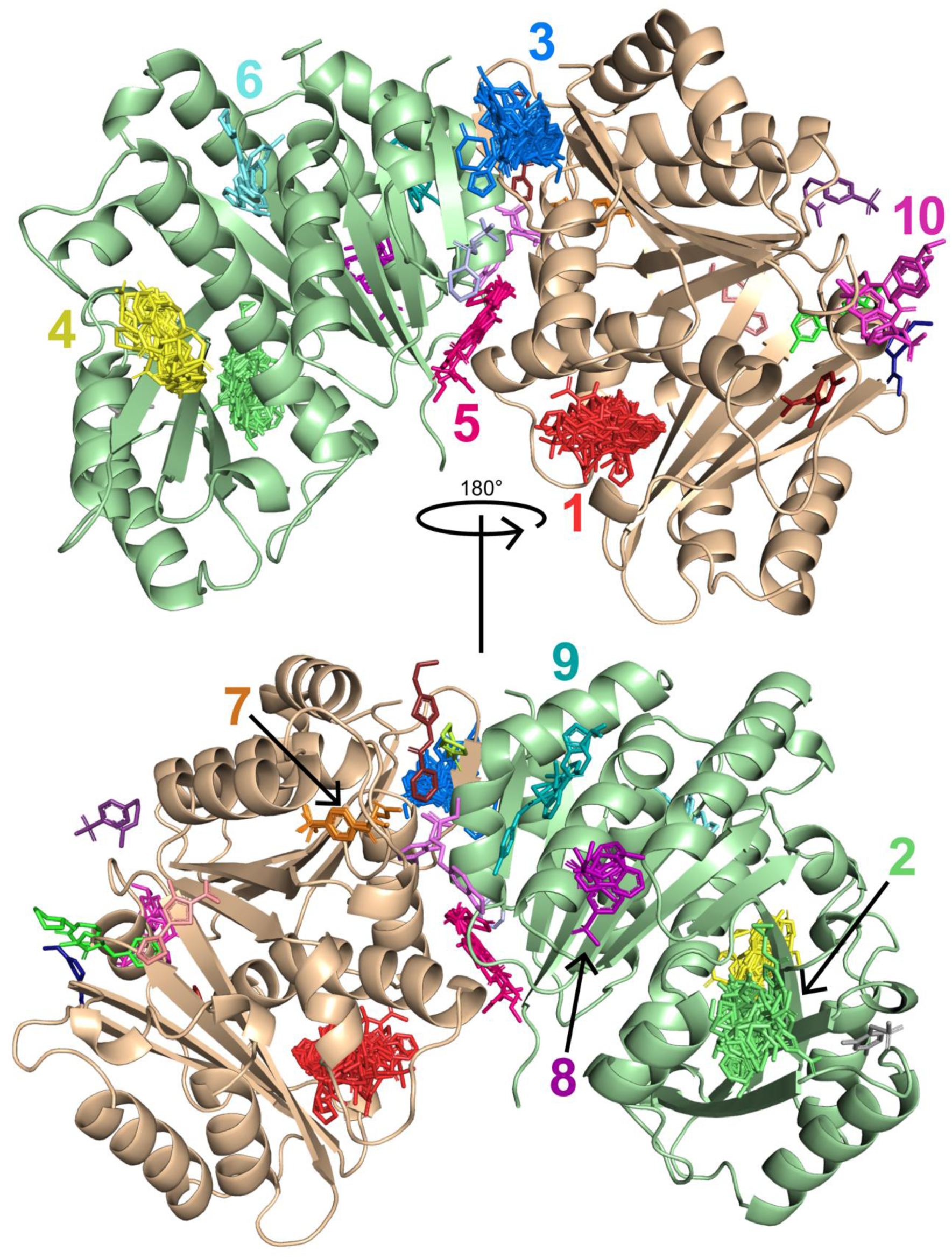
Fragments are colored according to site ID. The ten most populated fragment clusters are labeled; sites 11-20 are shown in Fig. S3. Chain A (ON protomer) is shown in pale green and chain B (OFF protomer) in wheat. Site assignments and structural context are summarized in Table 1.

**Table 1.** Summary of MtbFtsZ fragment-binding sites identified by crystallographic fragment screening. The number of unique fragments represents the number of chemically distinct fragments observed at each site; multiple binding poses of the same fragment within a site are counted once (*number of non-unique fragment poses in the site). Sites 1-10 are shown in Fig.2, and sites 11-20 are shown in Fig. S3. Previously identified sites include the ON and OFF nucleotide binding sites and the two cryptic pockets described in earlier MtbFtsZ structures.

| Site | Chain / state | No. of unique fragments | Site description | Previously identified | Structural / functional context |
| --- | --- | --- | --- | --- | --- |
| 1 | B / OFF | 59 | Cryptic pocket 2 | Yes | Dynamic pocket adjacent to H8; corresponding to site 4; strongly favors the OFF protomer |
| 2 | A / ON | 50 (52*) | H9-C-terminal beta-sheet pocket | No | State-dependent pocket; largely closed in chain B; overlaps a putative protofilament interface |
| 3 | A/B interface | 26 | Cryptic pocket 1 | Yes | Crystallographic dimer interface; formed by residues from both protomers, including sH2 |
| 4 | A / ON | 23 | Cryptic pocket 2 | Yes | Corresponding to site 1; substantially fewer fragments bind the ON protomer |
| 5 | A/B interface | 8 | Dimer-interface pocket | No | Adjacent to cryptic pocket 1 (site 3) |
| 6 | A / ON | 5 | Nucleotide-binding site | Yes | Fragments overlap the guanine-binding region; also at a putative protofilament interface |
| 7 | B / OFF | 4 | Nucleotide-binding site | Yes | OFF-state pocket; benzene-sulfonamide motif; near the Leu269 polymerization interface |
| 8 | A / ON | 4 | Fragment cluster | No | Putative polymerization-interface-proximal region |
| 9 | A / ON | 3 | Fragment cluster | No | Crystallographic dimer-interface region |
| 10 | B / OFF | 4 | T7-adjacent pocket | No | About 4 Å from site 18; polymerization-associated region |
| 11 | B / OFF | 1 | Single-populated site | No | Adjacent to the closed site 2 region in the OFF protomer |
| 12 | A/B interface | 1 | Dimer-interface site | No | Located between sites 3 and 5 |
| 13 | A/B interface | 2 | Interface-proximal site | No | Near Leu269-containing longitudinal interface; shares Asp84 interaction with site 20 |
| 14 | B / OFF | 2 | Low-populated site | No | Adjacent to the closed site 2 region in the OFF protomer |
| 15 | B / OFF | 1 | Single-populated site | No | Adjacent to the closed site 2 region in the OFF protomer |
| 16 | B / OFF | 1 | Single-populated site | No | Peripheral surface-binding site |
| 17 | A/B interface | 1 | Interface-proximal site | No | Near the Leu269-containing longitudinal polymerization interface |
| 18 | B / OFF | 1 | T7-loop-adjacent site | No | Z100642432 binds adjacent to T7; about 4 Å from site 10 |
| 19 | A / ON | 1 | Single-populated site | No | Peripheral surface-binding site |
| 20 | A/B interface | 1 | Dimer-interface site | No | Shares an Asp84 interaction with site 13 |

The presence of two conformationally distinct MtbFtsZ protomers in the asymmetric unit allowed ligand binding to analogous structural regions to be compared between the ON and OFF states. Pronounced differences in the number and identity of the fragments bound to equivalent regions of the two protomers were observed, indicating that the conformational state of MtbFtsZ alters the accessibility and ligandability of several pockets.

### Fragments recognize both the ON and OFF conformations of the nucleotide-binding site

Fragments were observed in the nucleotide-binding pockets of both MtbFtsZ conformational states, although the geometry and chemical characteristics of the two sites differed substantially (Figs. 3 and 4).

**Figure 3.**
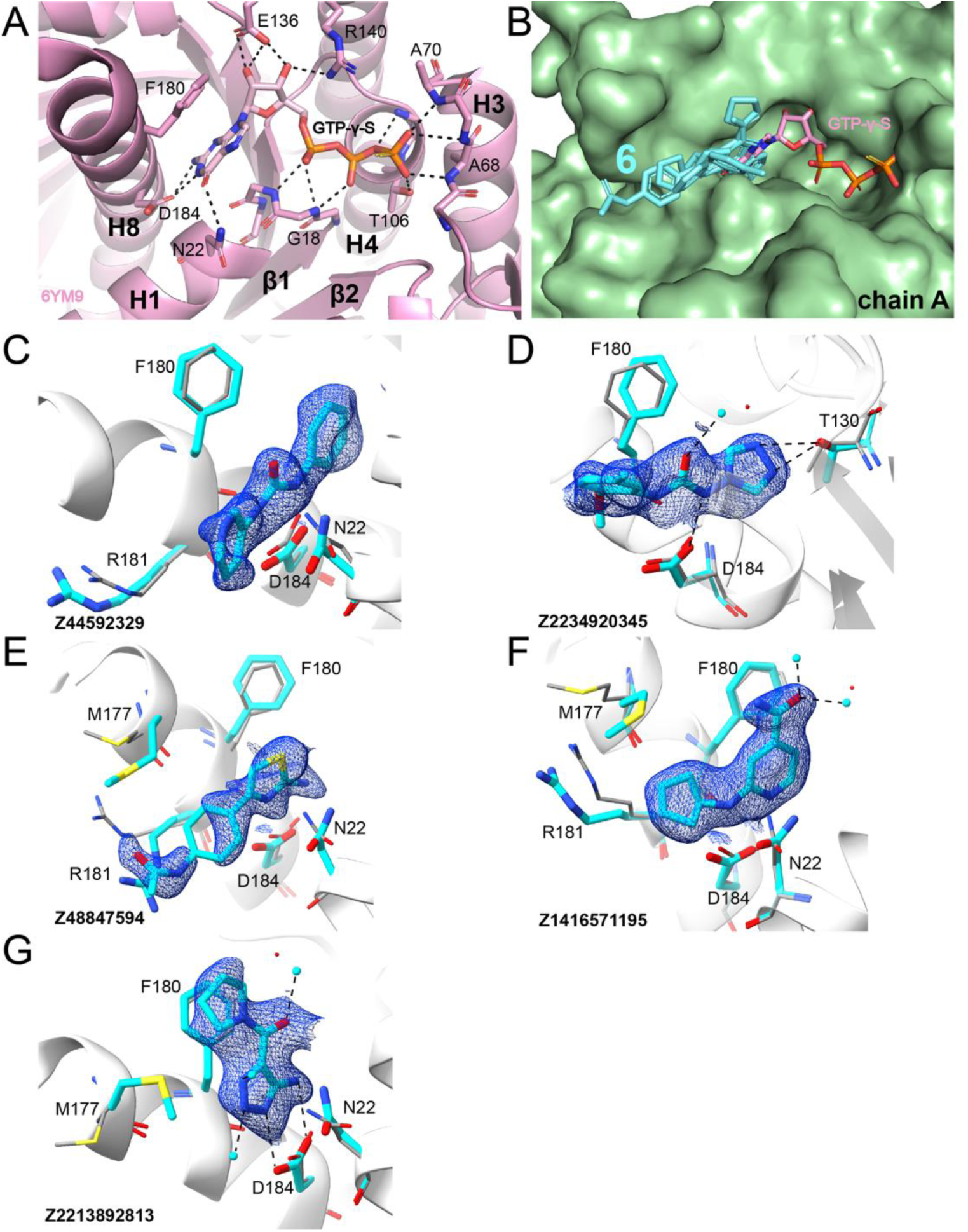
Fragments bind the ON-state nucleotide-binding pocket of MtbFtsZ. **A**) The binding site of GTPγ-S in FtsZ (pdb_00006ym9). Interactions between the protein and GTPγ-S are shown in black dashes. **B**) Surface representation of the ground state model (pdb_00015aa). GTPγ-S is superposed onto the structure. Fragments from site 6 are shown in aquamarine sticks. **C-G**) Fragments in the ‘ON’ GTP binding site are shown in stick representation. The ensemble model is represented as the changed state in aquamarine and the ground state (or atoms with a single conformation) shown in white/gray. Primary evidence of binding shown as PanDDA density event maps in blue mesh. Hydrogen bonding interactions are highlighted with black, dashed lines.

**Figure 4.**
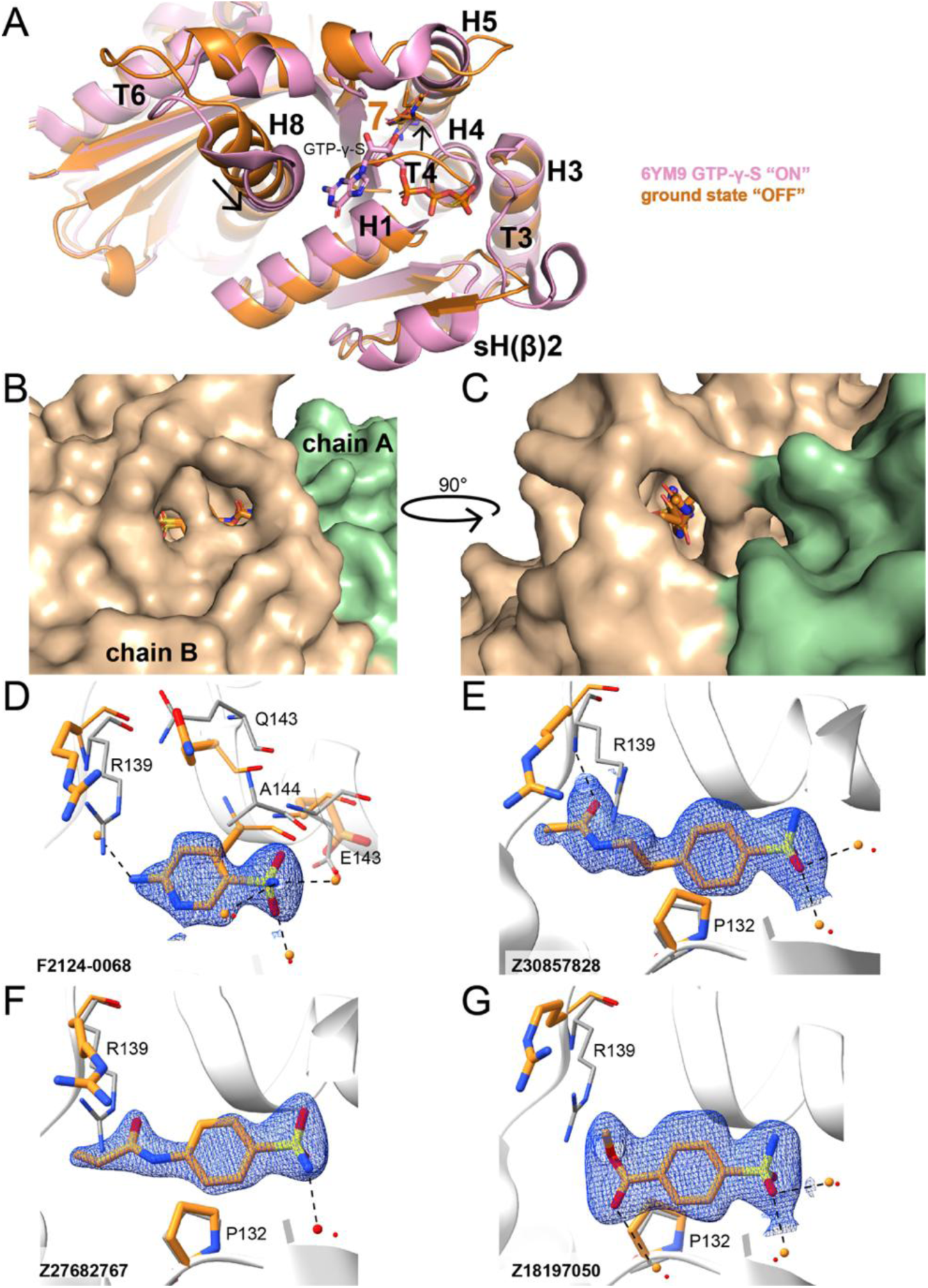
A conserved benzene-sulfonamide motif binds the OFF-state nucleotide-binding pocket. **A**) Superposition of ‘ON’ GTP-bound site (pdb_00006ym9) and ‘OFF’ GTP site (ground state model, this study). The ‘ON’ state is shown in pink. The ‘OFF’ state is shown in orange. Arrows indicate conformational changes required for GTP binding. **B-C**) Orthogonal surface representation of ground state model with site 7 fragments bound. Chain A (ON) is shown in pale green, chain B (OFF) is shown in tan. **D-G**) Fragments in the ‘OFF’ GTP binding site are shown in stick representation. The ensemble model is represented as the changed state in orange and the ground state (or atoms with a single conformation) shown in white/gray (or red in cases of water molecules). Primary evidence of binding shown as PanDDA density event maps in blue mesh. Hydrogen bonding interactions are highlighted with black, dashed lines.

Five fragments bind site 6, corresponding to the nucleotide-binding pocket of the ON protomer (Fig. 3). Superposition with GTPγ-S-bound MtbFtsZ shows that these fragments directly overlap the position occupied by the guanine moiety of the nucleotide (Fig. 3A-B). Fragment binding is supported by hydrogen-bonding and ionic interactions involving Thr130 and Asp184, while Met177, Phe180, Ala183, and Leu187 contribute hydrophobic contacts (Fig. 3C-G). Three fragments (Z2234920345, Z1416571195, and Z2213892813) additionally form hydrogen bonds with water molecules within the nucleotide binding pocket.

Four fragments bind site 7, corresponding to the nucleotide-binding region of the OFF protomer (Fig. 4). Relative to the ON state, bending of H8 is accompanied by disordering of the T3 loop and occupation of the nucleotide-binding region by the T4 loop, producing a substantially altered and smaller pocket (Fig. 4A-C). Despite the reduced size of the OFF-state pocket, four fragments were observed at this site. All four contain a pyridine-or benzene-sulfonamide group that adopts a similar orientation within the pocket (Fig. 4D-G). Few direct polar or electrostatic interactions support these poses; instead, the conserved aryl sulfonamide binding mode is primarily associated with hydrophobic interactions involving Pro132 and Ala144.

Thus, fragments can recognize both the ON state and the OFF state of MtbFtsZ. The distinct fragment chemotypes and interactions observed between these pockets further demonstrate that the two conformational states present chemically different ligand-binding environments.

### Conformational differences between MtbFtsZ protomers produce strongly asymmetric fragment binding

The most pronounced state-dependent fragment binding was observed at cryptic pockets 2 (sites 1 and 4) and at a previously uncharacterized pocket in the C-terminal domain (site 2; Fig. 5).

**Figure 5.**
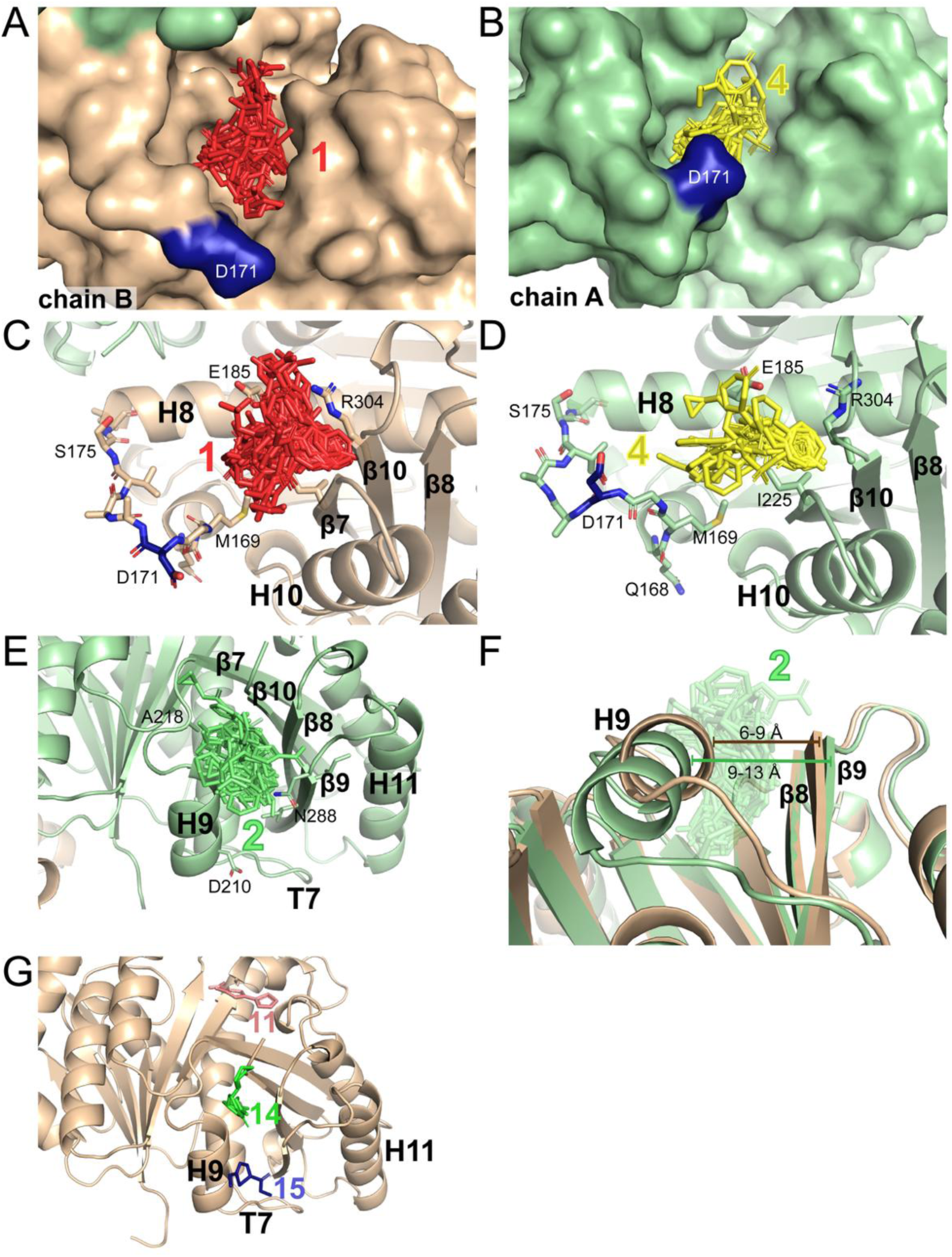
Conformational differences alter fragment binding at cryptic pocket 2 (sites 1 & 4) and site 2. **A-B**) Surface representations of cryptic pocket 2 in the OFF protomer (site 1; chain B, wheat) and ON protomer (site 4; chain A, pale green), respectively. Asp 171 is shown in blue. **C-D**) Fragments and residues forming sites 1 (**C**) and 4 (**D**). **E**) Fragments at site 2 in the ON protomer. **F**) Superposition of the C-terminal domains of the ON and OFF protomers illustrating displacement of H9 relative to β9 and closure of the corresponding site 2 pocket (sites 11, 14, 15) in the OFF protomer. **G**) Nearby fragment-binding sites 11, 14, and 15 in the OFF protomer (chain B).

Cryptic pocket 2 was the most highly populated region identified in the screen, containing 76 unique fragments across the two protomers. However, fragment binding was strongly asymmetric: 59 fragments bind site 1 in the OFF state of chain B, whereas 23 bind the corresponding site 4 ON state in chain A, with only six fragments shared between the two sites (Fig. 5A-D). The pocket is located adjacent to H8 and is formed in part by residues Ile225 and Arg304, which provide hydrophobic and hydrogen-bonding or ionic interactions, respectively. Additional pocket residues include Leu166, Met169, Glu185, Val186, Asn189, and Ser227.

Comparison of the two protomers provides a structural explanation for the difference in ligandability. Bending of H8 alters the position of the adjacent loop, highlighted by the displacement of Asp171 in Figures 5A-D, and changes the accessibility of the binding pocket. The substantially different number of fragments observed in sites 1 and 4 (cryptic pocket 2) therefore correlates with conformational differences between the two MtbFtsZ states. These observations are consistent with the previously reported conformational dependence of cryptic pocket 2, although the present screen demonstrates that some fragments can also bind the corresponding pocket in the ON state.

An even more pronounced asymmetry was observed at site 2, a previously uncharacterized pocket located between H9 and the C-terminal β-sheet (Fig. 5E-G). Fifty-two fragments bind this pocket in the ON protomer (chain A), making it the second most populated fragment cluster identified in the screen. Superposition of the C-terminal domains shows a 3-5 Å displacement that brings H9 closer to β9 in the OFF protomer (chain B), effectively closing the site 2 pocket (Fig. 5F). Only four fragments (sites 11, 14, and 15) occupy the corresponding site 2 pocket of the OFF protomer (chain B; Fig. 5G). The pocket is predominantly hydrophobic, with Ile214, Met215, Ala218, Leu258, Ile290, Ile308, and Ala310 frequently contacting bound fragments.

Together, sites 1, 4 and 2 demonstrate that relatively small conformational differences between the ON and OFF MtbFtsZ protomers can substantially alter pocket accessibility and fragment binding.

### Multiple fragment-binding sites occur along the crystallographic dimer interface

Several fragment-binding sites were identified along the interface between the two MtbFtsZ protomers in the asymmetric unit (Fig. 6; Table 1). The largest of these, site 3, corresponds to cryptic pocket 1 and contains 26 unique fragments (Fig. 6A, C-E).

**Figure 6.**
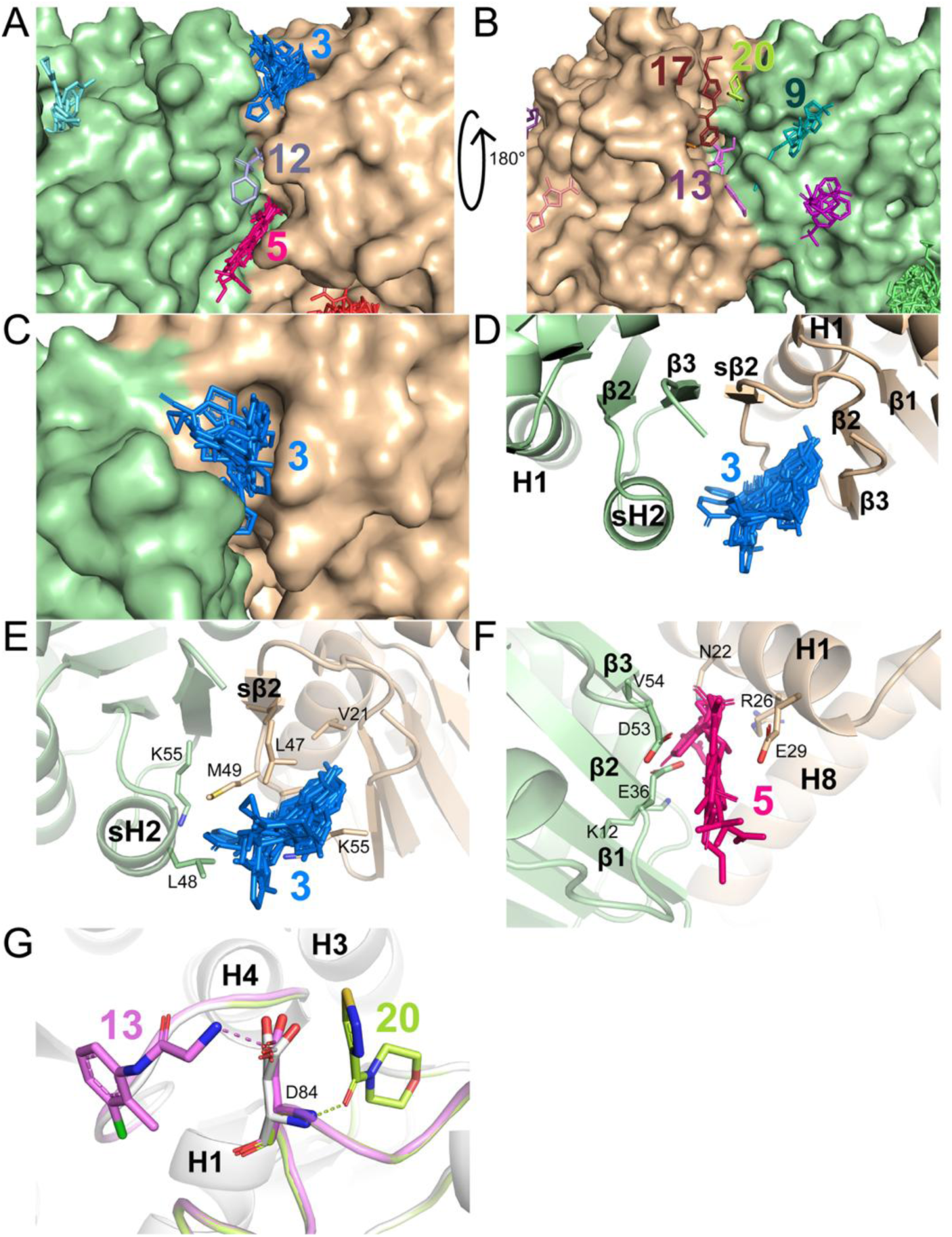
Fragments bind to the crystallographic dimer interface of FtsZ. **A-B**) The ground state model represented with a surface. Fragments are colored by site ID. Sites found at the crystallographic dimeric interface are labeled. Figures A and B are related by a 180° rotation. **C**) Surface representation of the ground state model with site 3 (cryptic pocket 1) fragments shown in blue sticks. **D**) Cartoon representation of the ground state model and secondary structure elements that form cryptic pocket 1 are highlighted. **E**) Key residues that form cryptic pocket 1 are shown with side chains in stick representation. **F**) Site 5 fragments shown in magenta with key residues that form the binding pocket shown in sticks from the ground state model. **G**) Sites 13 and 20 at the opposite face of the dimer-interface groove; Asp84, which contributes interactions to both sites, is shown in stick representation.

The site 3 pocket is formed by structural elements from both protomers (Fig. 6C), including H1, sH2/sβ2, β2, and β3. Relative to sH2 in chain A, chain B is rotated by ∼90°, positioning sβ2, and β3 adjacent to the binding pocket (Fig. 6D). Fragment binding is supported by hydrophobic contacts involving Ala44 and Leu48 from chain A and Val21, Ala39, Leu47, and Met49 from chain B while Ser50 and Lys55 provide additional hydrogen-bonding interactions (Fig 6E).

Several smaller sites extend along the same crystallographic interface. Site 5 contains eight fragments and is positioned adjacent to site 3, with contacts involving Lys12, Val54, Asn22, and Arg26 (Fig. 6F). Additional sites, 13, 17, and 20, occur along the opposite face of the dimer-interface groove (Fig. 6A-B). Sites 13 and 20 share interactions with Asp84 (Fig. 6G). Collectively, these observations reveal multiple ligandable regions distributed along the crystallographic MtbFtsZ dimer interface.

### Fragment-binding sites overlap putative FtsZ polymerization interfaces

Several fragment-binding sites are located at or near structural elements involved in FtsZ protofilament formation (Fig. 7). A fragment, Z100642432, binds adjacent to the T7 loop in chain B (site 18; Fig. 7A). Although no prominent direct interactions between the fragment and T7 loop residues occur within 4 Å, a cluster of four fragments at site 10 is located ∼ 4 Å from Z100642432 (Fig. 7B). The proximity of these sites defines an extended ligandable region adjacent to the T7 loop.

**Figure 7.**
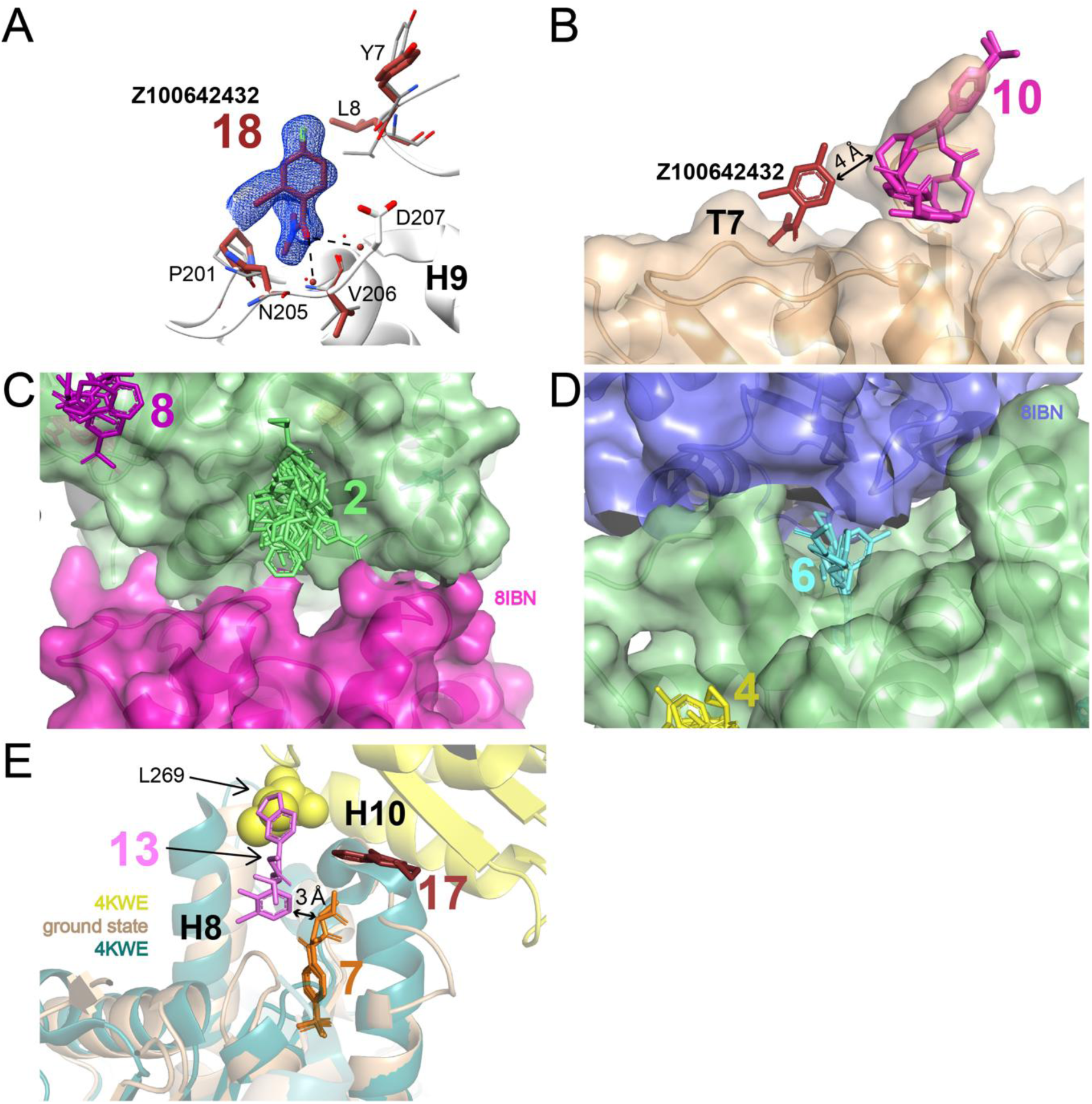
Fragment-binding sites overlap or lie near putative FtsZ protofilament interfaces. **A**) Site 18 fragment shown in sticks. Residues that form the binding pocket are shown in sticks and labeled. Primary evidence of binding shown as PanDDA density event maps in blue mesh. Hydrogen bonding interactions are highlighted with black, dashed lines. **B**) Site 10 fragments located ∼4 Å from site 18 shown in sticks, magenta. **C-D**) Sites that could disrupt polymerization interactions are shown in sticks. Chain A of the ground state model is shown in pale green surface. Hypothesized binding modes for FtsZ polymerization are reflected with KpFtsZ (pdb_00008ibn). Adjacent KpFtsZ protomers are shown in purple and magenta. **E**) Superposition of the MtbFtsZ ground-state model with the curved MtbFtsZ protofilament (pdb_00004kwe). Leu269 at the longitudinal interface is shown as spheres, and nearby fragment-binding sites 7, 13, and 17 are shown as sticks.

To identify additional fragment-binding sites positioned at potential longitudinal FtsZ interfaces, the MtbFtsZ structure was compared with available straight and curved FtsZ protofilament structures (Fig. 1C-D). Superposition with the straight *Klebsiella pneumoniae* FtsZ (KpFtsZ) protofilament places sites 2 and 6 at regions corresponding to interprotomer interfaces (Fig. 7C-D). Although MtbFtsZ and KpFtsZ have ∼50% sequence identity and individual interprotomer contacts may differ, the structural superposition places both fragment-binding sites at the protofilament interface.

The fragment-binding map was also compared with the previously determined curved MtbFtsZ protofilament structure (pdb_00004kwe). Leu269 forms a key component of the longitudinal interface in this structure. Sites 7, 13, and 17 are located near this Leu269-containing interface (Fig. 7E). Thus, multiple fragment-binding sites identified in the screen overlap or lie proximal to structural regions implicated in FtsZ protofilament assembly.

### Protein and water interactions differ substantially among fragment-binding sites

Protein-fragment and water-mediated interactions were compared across the binding sites to characterize the chemical environments sampled by the fragment screen (Fig. 8).

**Figure 8.**
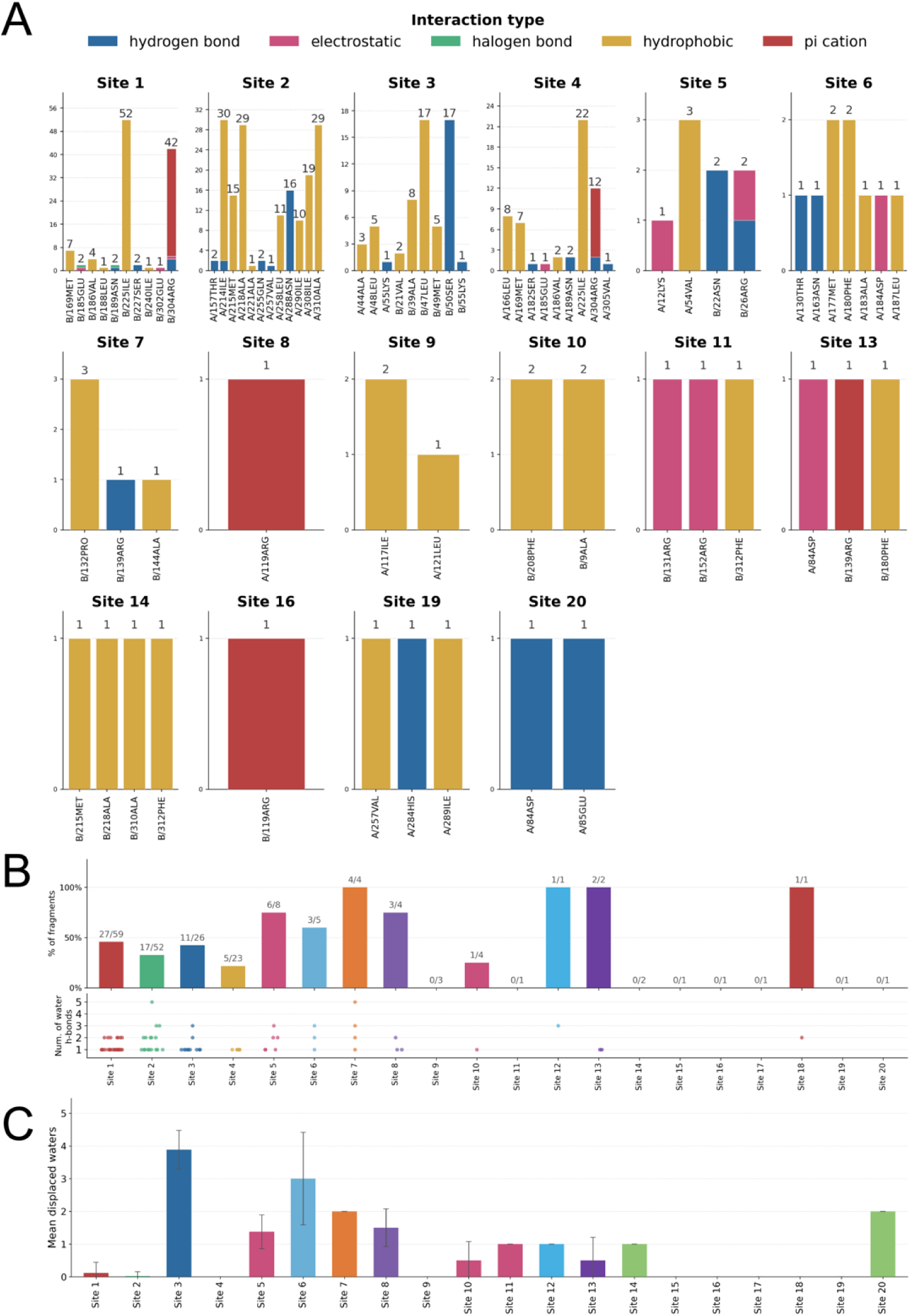
Fragment interactions with FtsZ and water molecules by site. **A**) Each bar graph represents a different fragment binding site. The colors of the bar represent the type of interaction (blue: hydrogen bond, pink: ionic, green: halogen bond, yellow: hydrophobic, red: pi cation). **B**) The number of fragments within each site that form at least one hydrogen bond with water are shown in top bar graph. The jitter plot shows the number of water molecules per fragment (e.g. a point at 4 means that one fragment hydrogen bonds with 4 water molecules). **C**) Average number of ground-state water molecules displaced per unique fragment at each binding site.

The highly populated sites 1-4 contained the largest numbers of protein-fragment interactions, although the relative contributions of hydrophobic, hydrogen-bonding, electrostatic, and other contacts differed among sites (Fig. 8A). The previously uncharacterized site 2 was dominated by hydrophobic interactions, consistent with the composition of the pocket. Sites 1 and 4, corresponding to cryptic pocket 2 in the two conformational states, also displayed different interaction profiles despite being formed by analogous regions of the protein.

Water-mediated interactions also varied among sites (Fig. 8B). Approximately half of the fragments occupying site 1 formed at least one hydrogen bond with a nearby water molecule, whereas approximately one-fifth of fragments in site 4 formed such an interaction. Seventeen fragments at site 2 interacted with 1 to 5 water molecules. In contrast, relatively few fragments in several of the lower populated sites formed water-mediated interactions.

Comparison with the PanDDA ground-state model further identified differences in water displacement upon fragment binding (Fig. 8C). Fragments occupying the ON nucleotide-binding pocket displaced ∼3 waters on average, whereas fragments in the OFF nucleotide-binding site displaced ∼2. Site 3 displayed the greatest water displacement, with approximately four ground-state water molecules displaced per unique fragment pose. In contrast, fragment binding to sites 1 and 4 resulted in little average water displacement.

These results show that the identified pockets differ not only in fragment occupancy and conformational accessibility but also in the balance of direct protein contacts, water-mediated interactions, and displacement of ordered solvent.

### Chemical similarity analysis identifies a recurrent benzimidazole scaffold in the cryptic pockets

Pairwise chemical similarity among the crystallographic hits was evaluated using Tanimoto coefficients and maximum common substructure (MCS) analysis (Fig. 9). The majority of the fragment pairs had Tanimoto coefficients below 0.4, indicating that the overall hit set is chemically diverse (Fig. 9A). The MCS distribution showed a prominent maximum at six shared heavy atoms, consistent with the frequent occurrence of aromatic six-membered rings among the fragments (Fig. 9B).

**Figure 9.**
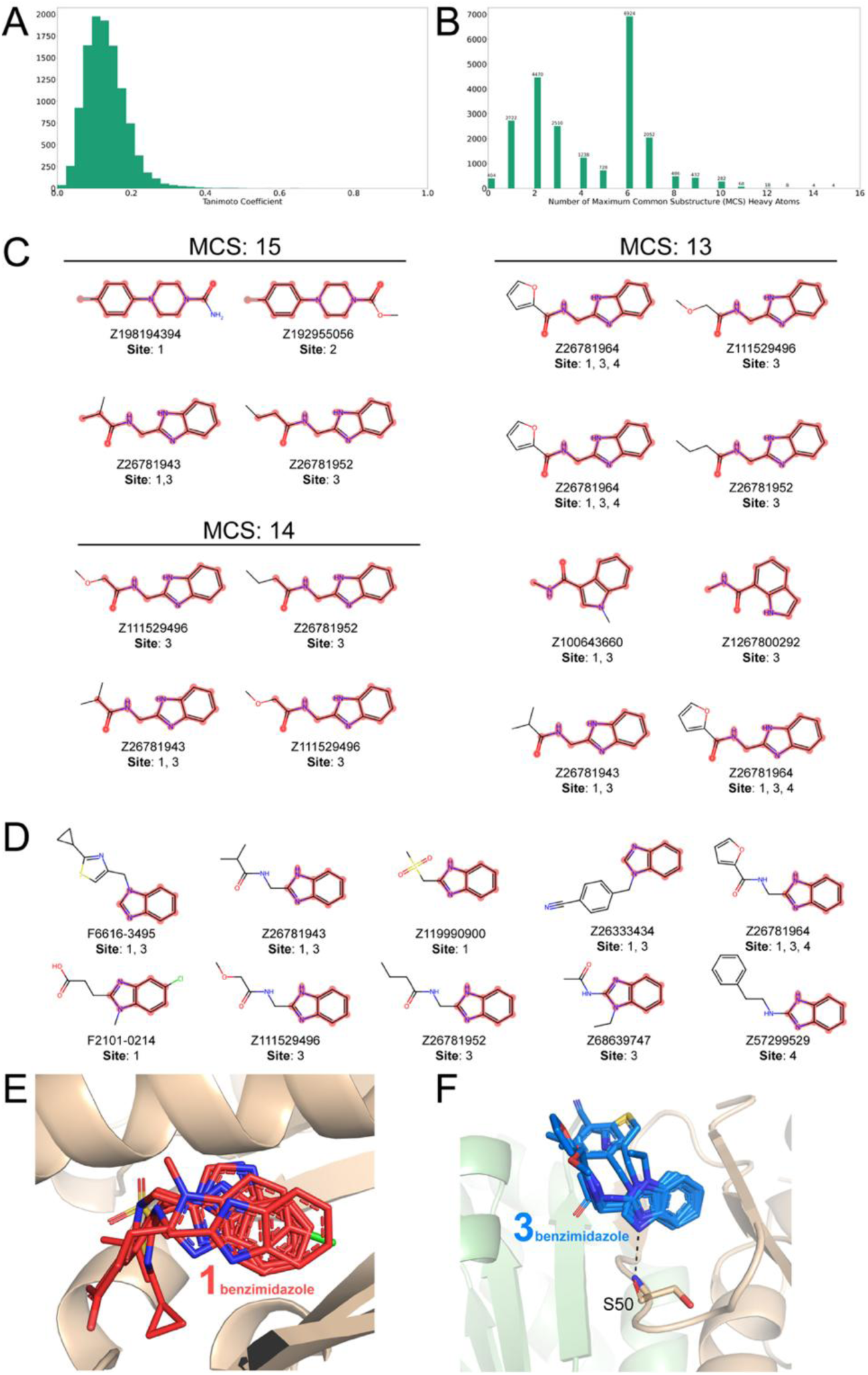
Chemical similarity analysis identifies recurrent scaffolds among MtbFtsZ fragment hits. **A**) Distribution of pairwise Tanimoto coefficients among fragment hits. **B**) Distribution of maximum common substructure (MCS) size, expressed as the number of shared heavy atoms. **C**) Fragment pairs with the largest MCS values. **D**) Fragment hits containing the recurrent benzimidazole or related bicyclic scaffold. **E-F**) Superposition of benzimidazole-containing fragments bound at sites 1 (**E**) and 3 (**F**). Ser50, which forms a conserved hydrogen bond with the benzimidazole group at site 3, is shown in stick representation.

Despite the overall chemical diversity of the hit set, analysis of fragment pairs with the largest common substructures revealed a recurrent scaffold. The largest MCS identified contained 15 heavy atoms corresponding to a 1-(4-fluorophenyl)piperazine group (Fig. 9C). Among fragment pairs sharing at least 13 heavy atoms, six of the seven highest-ranking fragments contained a benzimidazole or related bicyclic group.

Ten crystallographic hits contain this scaffold, and all bind to previously characterized coumarin-binding pockets (Fig. 9D-F). Six bind site 1, where the orientation of the benzimidazole group varies among each unique fragment (Fig. 9E). Seven bind site 3, where the benzimidazole groups adopt a more strongly conserved orientation (Fig. 9F). In site 3, a conserved hydrogen bond is observed between the backbone nitrogen of Ser50 and a nitrogen atom of the benzimidzaole group.

Thus, although the fragment screen sampled a chemically diverse collection of ligands, recurrent chemical features emerged at specific MtbFtsZ pockets, with benzimidazole-containing fragments preferentially recognizing the previously described cryptic coumarin-binding sites.

### Fragment binding was not observed or could not be assessed at additional regions of interest

No fragments were observed at the T9 loop or the associated protomer interface. MtbFtsZ structure (pdb_00005v68) identified an unusual intermolecular interaction in which the T9 loop of one protomer occupies the nucleotide-binding site of an adjacent protomer^14^. The intrinsically disordered C-terminal region of MtbFtsZ (residues 313-379) was not resolved in the structures obtained in this study; therefore, potential fragment binding to this region could not be assessed by crystallography. In addition, no fragments were observed directly within the previously characterized SaFtsZ allosteric pocket when mapped onto the corresponding region of MtbFtsZ (Fig. S4).

## Discussion

The crystallographic fragment screen presented here substantially expands the known ligand-binding landscape of MtbFtsZ. Fragments were observed across the nucleotide-binding sites, previously characterized cryptic pockets, the crystallographic dimer interface, and regions associated with protofilament assembly. Of the 20 identified binding sites, 15 have not been previously described as ligand-binding sites in MtbFtsZ. These results demonstrate that MtbFtsZ contains a substantially broader range of ligandable surfaces than indicated by the relatively limited number of previously reported inhibitor-bound structures. Rather than identifying a single dominant drug-binding pocket, the screen reveals a distributed and conformationally dynamic ligand-binding landscape that provides multiple starting points for the development of chemical probes and, potentially, inhibitors of MtbFtsZ function.

A notable feature of the screen is the pronounced dependence of fragment binding on the conformational state of MtbFtsZ. The crystallographic asymmetric unit contains two protomers corresponding to the previously described ON and OFF switch configurations, allowing ligand binding to structurally analogous regions to be compared under identical fragment-soaking conditions. Despite their similar overall folds, several equivalent regions displayed markedly different fragment-binding profiles. The strongest example is cryptic pocket 2, where 59 fragments bind site 1 in the OFF protomer compared with 23 at the corresponding site 4 in the ON protomer, with only six fragments shared between the two sites. A similarly pronounced conformational dependence was observed at site 2, where 52 fragments occupy a pocket between H9 and the C-terminal β-sheet in the ON protomer, while structural rearrangement substantially reduces accessibility to the corresponding region in the OFF protomer. These observations demonstrate that relatively small conformational rearrangements in MtbFtsZ can substantially alter its ligand-binding landscape. Molecules that preferentially recognize one of these conformational states could therefore provide useful chemical probes for investigating the relationship between MtbFtsZ conformation and function and, following further optimization and validation, may provide a strategy for stabilizing different MtbFtsZ conformational states.

The extensive fragment binding observed within the previously identified cryptic pockets further supports these regions as ligandable hotspots in MtbFtsZ. Alnami *et al.* identified two cryptic pockets using structures containing 4-hydroxycoumarin, with pocket formation linked to conformational changes within MtbFtsZ. In the present unbiased fragment screen, cryptic pocket 2 (sites 1 and 4) was the most highly populated region of the protein, while cryptic pocket 1 (site 3) contained 26 unique fragments. The ability of chemically diverse fragments to occupy these pockets indicates that their ligandability is not restricted to coumarin derivatives. Chemical similarity analysis further identified a recurrent benzimidazole or related bicyclic scaffold among fragments occupying MtbFtsZ cryptic pockets. Ten hits containing this scaffold were identified, with strong structural conservation of the benzimidazole binding orientation at site 3, including a conserved hydrogen-bond interaction with the backbone nitrogen of Ser50. These recurring structural and chemical features provide potential starting points for fragment growth and structure-guided optimization. However, although the previously reported coumarin structures and the present fragment screen establish that these pockets can accommodate small molecules, the functional consequences of occupying these sites remain to be established experimentally.

Fragments were also observed at or near several regions directly associated with nucleotide binding and protofilament assembly. Five fragments occupy the nucleotide-binding pocket of the ON protomer and overlap the guanine-binding region, demonstrating that this highly conserved functional site can accommodate non-nucleotide fragment chemotypes. Four additional fragments occupy the structurally reorganized nucleotide-binding region of the OFF protomer and share a conserved aryl sulfonamide group despite having different substituents. The ability to recognize both nucleotide-site conformations raises the possibility of developing ligands selective for distinct states of the MtbFtsZ nucleotide-binding pocket. Additional fragments occur near the T7 loop and longitudinal protofilament interfaces. In particular, site 18 is adjacent to T7 and lies ∼4 Å from the site 10 fragment cluster, defining an extended ligandable region that could be explored by fragment growing or linking. Similarly, sites 7, 13, and 17 lie near the Leu269-containing longitudinal interface observed in the curved MtbFtsZ protofilament. These structural relationships provide experimentally testable starting points for targeting regions involved in FtsZ assembly, although the present crystallographic data alone do not establish that occupation of these sites disrupts polymerization or GTPase activity.

*Staphylococcus aureus* FtsZ inhibitors such as TXH9179 bind a hydrophobic pocket located between the core helix and the C-terminal β-sheet^23,24^. Structural comparison of SaFtsZ with MtbFtsZ shows that the C-terminal β-sheet of MtbFtsZ is positioned ∼10 Å closer to the core helix (H8), substantially reducing the volume and accessibility of the corresponding allosteric pocket (Fig. S4). Consistent with this structural difference, no fragments were observed directly within the canonical SaFtsZ allosteric pocket in the present screen. However, several ligandable sites were identified in the surrounding region. In the ON protomer, sites 2 and 4 are located within ∼10 Å of the SaFtsZ allosteric pocket, whereas sites 1,14, 15, and 18 occupy nearby regions when SaFtsZ is superimposed onto the OFF protomer (Fig. S4). Thus, although the canonical SaFtsZ allosteric pocket does not appear to be directly accessible in the ON/OFF MtbFtsZ conformations observed in the present study, the surrounding region contains alternative ligandable pockets that may provide opportunities to perturb this conformationally important region. Determining whether ligand binding at these sites influences MtbFtsZ conformation or function will require further biochemical investigation.

Other regions of MtbFtsZ were either not occupied by fragments or could not be evaluated in the present crystallographic screen. Lazo *et al.* previously reported an unusual intermolecular interaction in which the T9 loop of one MtbFtsZ protomer occupied the nucleotide-binding pocket of an adjacent protomer^14^. The biological significance of this interaction remains uncertain, and no fragments were observed at the T9 loop or its associated protomer interface. The intrinsically disordered C-terminal region of MtbFtsZ also represents an important limitation of the crystallographic analysis. This region participates in interactions with components of the divisome, including SepF and FtsW, with residues Asp367-Asp370 implicated in the interaction with FtsW^10,11^. However, residues 313-379 were not resolved in the structures obtained here. Consequently, the absence of crystallographic fragment hits cannot be interpreted as evidence that this region is not ligandable; potential fragment interactions with the disordered C-terminal regions simply could not be assessed with our structures.

Several additional limitations should be considered when interpreting the fragment-binding sites identified in this study. Crystallographic fragment screening establishes that a small molecule can engage a specific protein surface or pocket but does not itself establish binding affinity or functional modulation. The fragments identified here are expected to represent starting points, and quantitative binding affinities and effects on MtbFtsZ polymerization or GTPase activity were not measured as part of this screen. Furthermore, accessibility of individual pockets may be influenced by the conformational states and intermolecular contacts present within this hexagonal crystal form. Sites located at the crystallographic dimer interface therefore require additional validation to establish whether equivalent ligandable conformations are populated in solution. These considerations are important when assigning potential functional significance to newly identified sites. Quantitative binding measurements, MtbFtsZ polymerization and GTPase assays, and structure-guided fragment optimization will be necessary to determine which of these sites can be exploited to modulate MtbFtsZ function.

Together, these results provide a structural framework for exploring MtbFtsZ beyond the small number of ligand-binding sites characterized previously. The identification of state-dependent pockets, recurrent chemical scaffolds, and ligandable regions associated with nucleotide binding and protofilament interfaces establishes multiple avenues for subsequent fragment optimization and functional testing. More broadly, the pronounced differences in fragment binding between the ON and OFF conformations highlight the importance of protein conformational state in defining the ligandability of MtbFtsZ. Converting these crystallographic hits into higher-affinity molecules and determining their effects on MtbFtsZ assembly and GTPase activity will be important next steps toward establishing which of these newly identified pockets can be exploited for chemical modulation of this essential bacterial cell-division protein.

## Methods

### Expression and purification of FtsZ

FtsZ was expressed and purified by Genscript. *FtsZ* was codon optimized, cloned into the pET-28b(+) vector, and expressed in BL21 Star^TM^ (DE3). The plasmid sequence is available in Supplemental File 1. Expression was scaled up to 2L for subsequent purification. FtsZ was purified as described previously using thrombin cleavage of an N-terminal His-tag^14^. Size exclusion chromatography was used as the final step in purification into the storage and crystallization buffer of 50 mM Tris pH 7.8, 200 mM NaCl, and 100 mM KCl. The protein was concentrated to 2.19 mg/ml and flash frozen in liquid nitrogen.

### Crystallization of FtsZ

FtsZ was screened for crystallization conditions using the Index HT screen (Hampton Research). Initial hits were discovered in F6 (0.2 M Ammonium sulfate, 0.1 M BIS-TRIS pH 5.5, 25% (w/v) PEG 3,350). A grid search of nearby concentrations led to the final condition of 0.1 M ammonium sulfate, 15.6% (w/v) PEG 3350, 0.1 M Bis-Tris pH 5.0. Crystal trays optimized for fragment screening were formed using an Oryx8 (Douglas Instruments) in 288-drop SWISSCI MRC 3-well plates with 0.3 µl FtsZ, 0.2 µl reservoir solution, and 0.1 µl microseeds.

### Fragment screening

Fragment screening was performed at the NSLS-II fragment screening facility^34^. Crystal trays were imaged using the Crystal Shifter (Oxford Lab Technologies) to ensure that only drops with crystals were selected for soaking. An inhouse python application was used to track crystal soaking metadata in a SQLite database (https://github.com/NSLS2/mxplate). The fragment libraries screened included DSi-Poised library (DSIP; Enamine, Kyiv, UA) and Fragment Diversity Set III (FDS-III; Life Chemicals, Niagara-on-the-Lake, CA). The DSIP stock concentration was 500 mM and the FDS-III stock concentration was 100 mM. DMSO tolerance tests were performed at incremental volumes of DMSO and soaking times to find the highest DMSO concentration that could be used without impacting diffraction data quality. 30 µl (final concentration of approximately 10% v/v DMSO) of fragment solutions were transferred to crystal drops by acoustic ejection using an Echo 550 liquid handler (Beckman Labcyte). The average soak time was 152 ± 59 minutes (ranging from 67 to 305 minutes). Incubations of crystals with fragment solutions were performed at room temperature. Crystals were harvested with MiTeGen (Ithaca, NY, USA) microloops and a Crystal Shifter (Oxford Lab Technologies) followed by vitrification in liquid nitrogen.

### Data collection

X-ray diffraction data were collected at the AMX beamline (17-ID-1) at the National Synchrotron Light Source II at Brookhaven National Laboratory (Upton, NY). 843 datasets were obtained using the automated data collection workflow^32,35^. Diffraction data were indexed, integrated, and scaled with XDS^36^, merged with AIMLESS^37^, through autoPROC and fast_dp processing pipelines^38,39^. Diffraction datasets with resolution lower than 2.8 Å were excluded from subsequent Pan-Dataset Density Analysis (PanDDA).

### Data processing and model building

Pan-Dataset Density Analysis (PanDDA; installed in CCP4 v7.0.065) was used to identify potential binding events and model fragments^40^. Our current pipeline includes three runs of *pandda.analyse*, which parallels Approach 4 in Erlanson et al ^31^. First, datasets were auto-phased and refined using DIMPLE (CCP4) with 2Q1X (pdb_00002q1x) as a starting model^41^. The first *pandda.analyse* run was used to assess the hit rate and any conformational heterogeneity. AceDRG (CCP4) was used to generate the ligand CIF restraint files from the vendor-provided SMILES strings for each compound^42^.

The ground state model was generated in the second *pandda.analyse* run by calculating the highest resolution average map in *pandda.analyse* (with calculate_first_average_map_only set to True). The starting model used for DIMPLE in the previous *pandda.analyse* run was docked into the average map with Phenix’s Dock in map tool^43^. The model was manually inspected to best represent the average map. Finally, the ground state model was superimposed back to its starting position commensurate with the crystallographic origin. Using the *pandda.inspect* GUI, ligands were placed into appropriate event map density and changes were made to the local environment (5-10 Å) to represent any conformational changes to the protein and water molecules upon fragment binding^44^. The modeled ligand(s), along with any coordinate changes to the surrounding protein or water molecules, represent the changed state.

To refine occupancies of fragment binding events in each crystal, the ground state and changed state models were combined into an ensemble model. In our ensemble models, the changed state is represented with alt_id A and the ground state (or unchanged state) is represented with alt_id B. Occupancy groups were generated by spatially clustering all atoms with altloc labels using a periodic Euclidean distance function and the DBSCAN algorithm (*ԑ*=3.3 Å) as implemented in scipy. Occupancy groups are restrained such that all of the atoms found in the same altloc label within a DBSCAN-cluster have the same occupancy. The total occupancies of the ground state and the changed state within a cluster should sum to full occupancy if the chemical entity is present in both states. Ensemble models were refined with Refmac using external harmonic restraints enforced for each chain in the model (sigma=0.02)^45^. coordinates and associated structure factors for the fragment-bound MtbFtsZ structures were deposited in the Protein Data Bank (PDB) as a group deposition (G_1002379).

Each hit structure was deposited as an individual PDB entry. PanDDA event maps, which are the primary evidence used for ligand placement, are appended as additional *P*1 structure factor CIF blocks in hit structure factor CIF files. Remaining datasets for which no fragment ligand was modeled, but were used in PanDDA analysis, are included in the group deposition under a single entry (pdb_000015aa). The structure factors for each of these unmodeled datasets can be found as separate blocks within the structure factor CIF file of that entry. PDB accession codes, corresponding fragment identifiers, binding sites, and crystallographic statistics are provided in Supplementary Table S1. Fragment soaking metadata for both hit and unmodeled structures are included in the _diffrn.crystal_treatment key value pairs of structure factor CIF blocks. Occupancy group restraints and group deposition files were prepared with the NSLS-II fragflows software package (https://github.com/NSLS2/fragflows).

### Fragment clustering, interactions, and chemical similarity

To assign each fragment to a site, each ligand pose was identified. An anchor point was determined by finding ligand atoms within 4 Å of protein atoms. The centroid of the ligand atoms was calculated. The centroids of each ligand were compared so that any ligand within 4 Å were assigned to the same site. To ensure that overlapping ligands were also assigned the same site, despite the distance cutoff, single-linkage clustering was applied to the ligand anchors. The sites were manually inspected to consolidate any sites if necessary.

Protein-ligand and water-ligand interactions were calculated with gemmi and RDKit^46,47^. The following protein-ligand interactions were searched for: hydrogen bonds, ionic contacts, halogen bonds, hydrophobic contacts, pi-pi stacking, and pi-cation interactions. Distance and angle thresholds for all interaction types, besides hydrophobic, were relaxed based on the coordinate error associated with each structure. To identify displaced water molecules, water residue coordinates found only in the ground state model were compared to ligand coordinates. An ordered solvent molecule must be within 2.5 Å of the ligand to be displaced.

Tanimoto coefficients and maximum common substructure (MCS) pairs were calculated using RDKit. Pairwise chemical similarity was calculated iteratively for each hit molecule. Top MCS pairs were visualized using RDKit^47^.

## Supporting information

ftsz_supplemental_table1

pET-28b-plus_MtbFtsZ

## Data availability

Atomic coordinates (for hit structures), structure factor amplitudes (for all datasets), and PanDDA evidence maps (for hit structures) have been deposited in the Protein Data Bank under the Group Deposition accession code: G_1002379^48^. Analysis of PDB deposited structures are provided with this manuscript and supplemental information.

## Acknowledgements

The authors thank Jean Jakoncic for assistance with model building, data collection, and manuscript review. This research used resources 17-ID-1 of the National Synchrotron Light Source II, a U.S. Department of Energy (DOE) Office of Science User Facility operated for the DOE Office of Science by Brookhaven National Laboratory under Contract No. DE-SC0012704. The Center for BioMolecular Structure (CBMS) is primarily supported by the National Institutes of Health, National Institute of General Medical Sciences (NIGMS) through a Center Core P30 Grant (P30GM133893), and by the DOE Office of Biological and Environmental Research (KP1605010). The study was funded by by the National Institutes of Health P30 Grant (P30GM133893) to the Center for BioMolecular Structure at Brookhaven National Laboratory; by the Department of Energy (DOE) Office of Biological and Environmental Research (KP1607011). The funder played no role in study design, data collection, analysis and interpretation of data, or the writing of this manuscript.

## Author contributions

E.L. and D.K., conceptualization; E.L. and D.K., methodology; E.L. and D.K., investigation; D.K. and K.M., data curation; K.M., formal analysis; K.M., visualization; E.L. and K.M., writing-original draft; E.L, D.K., K.M., writing-review & editing; E.L. and D.K., funding acquisition.

## Abbreviations

TB: tuberculosis
MDR-TB: multidrug-resistant TB
(Mtb)FtsZ: (*Mycobacterium tuberculosis*) filamentous temperature-sensitive mutant Z
sH(β)2: switch helix (beta strand) 2
KpFtsZ: *Klebsiella pneumoniae* FtsZ
SaFtsZ: *Staphylococcus aureus* FtsZ
EcFtsZ: *Escherichia coli* FtsZ
AMX: Highly Automated Macromolecular Crystallography
DSIP: DSi-Poised library
FDS-III: Fragment Diversity Set III
PanDDA: pan-dataset density analysis
PDB: Protein Data Bank
MCS: maximum common substructure

## Supplemental Figures

**Figure S1.**
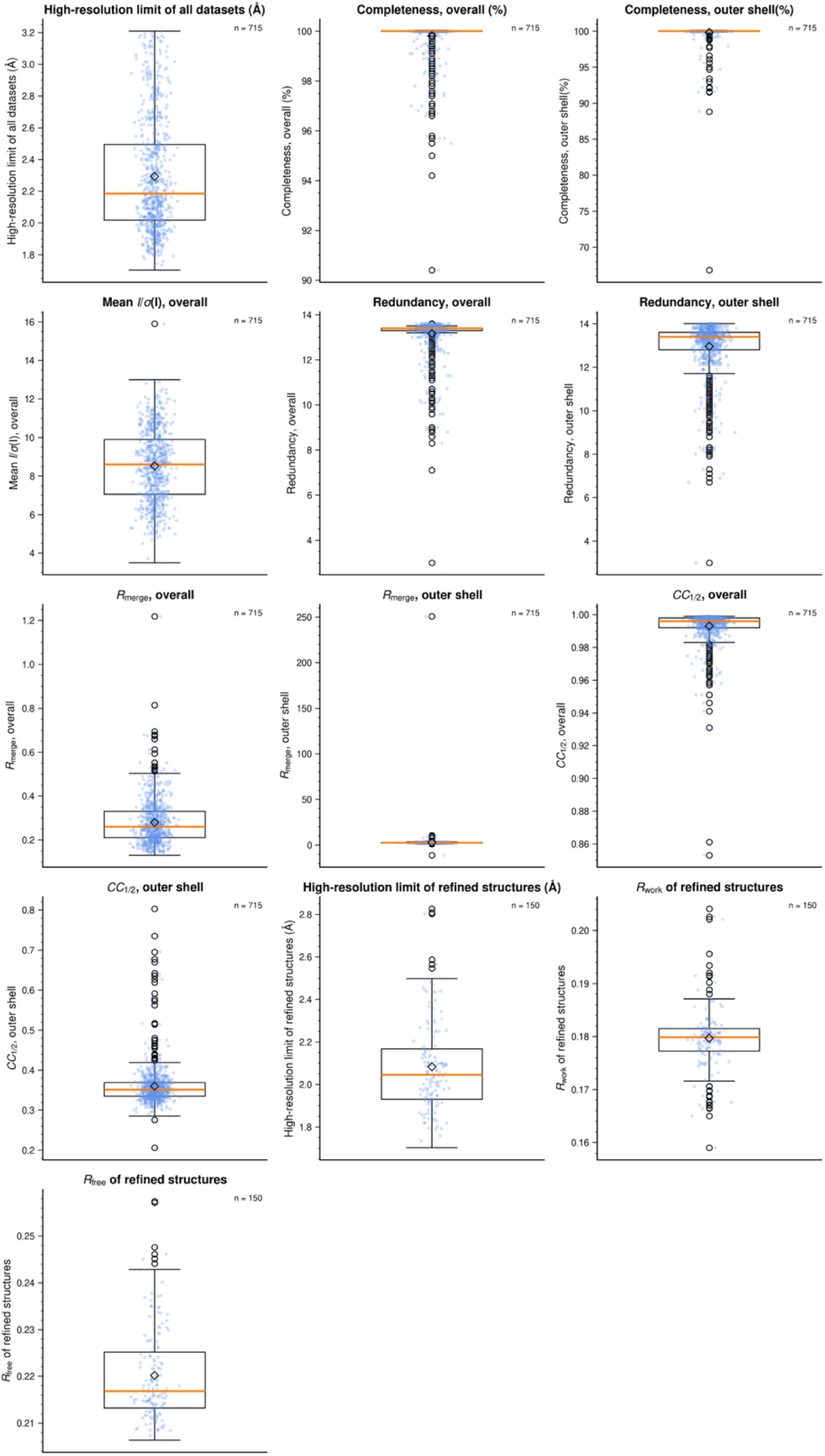
Data collection and refinement statistics shown as boxplots. The median for each parameter is represented by the orange line in the box. The mean is represented as the diamond. Outliers are shown in white circles.

**Figure S2.**
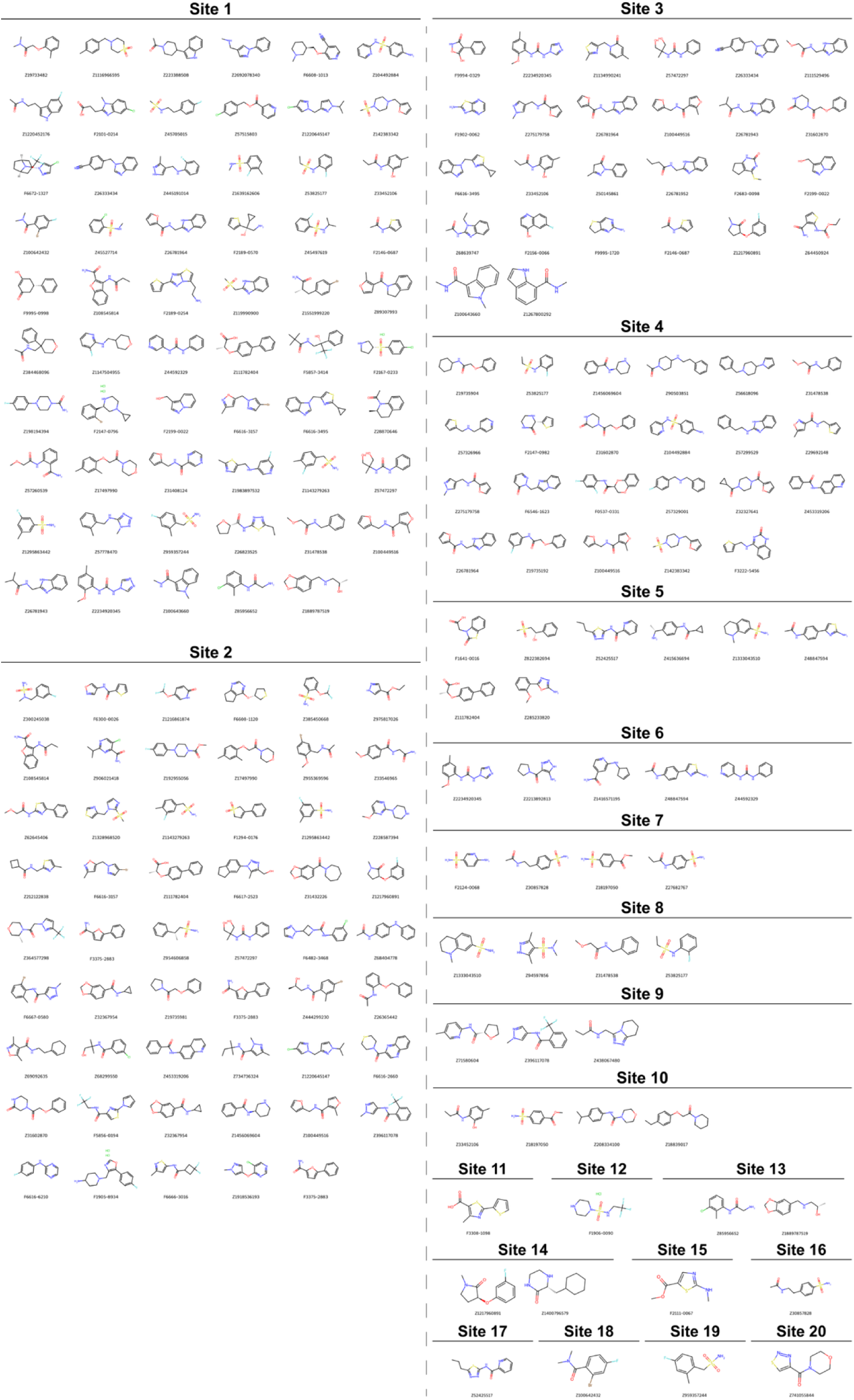
Chemical structure of hit fragments grouped by binding site.

**Figure S3.**
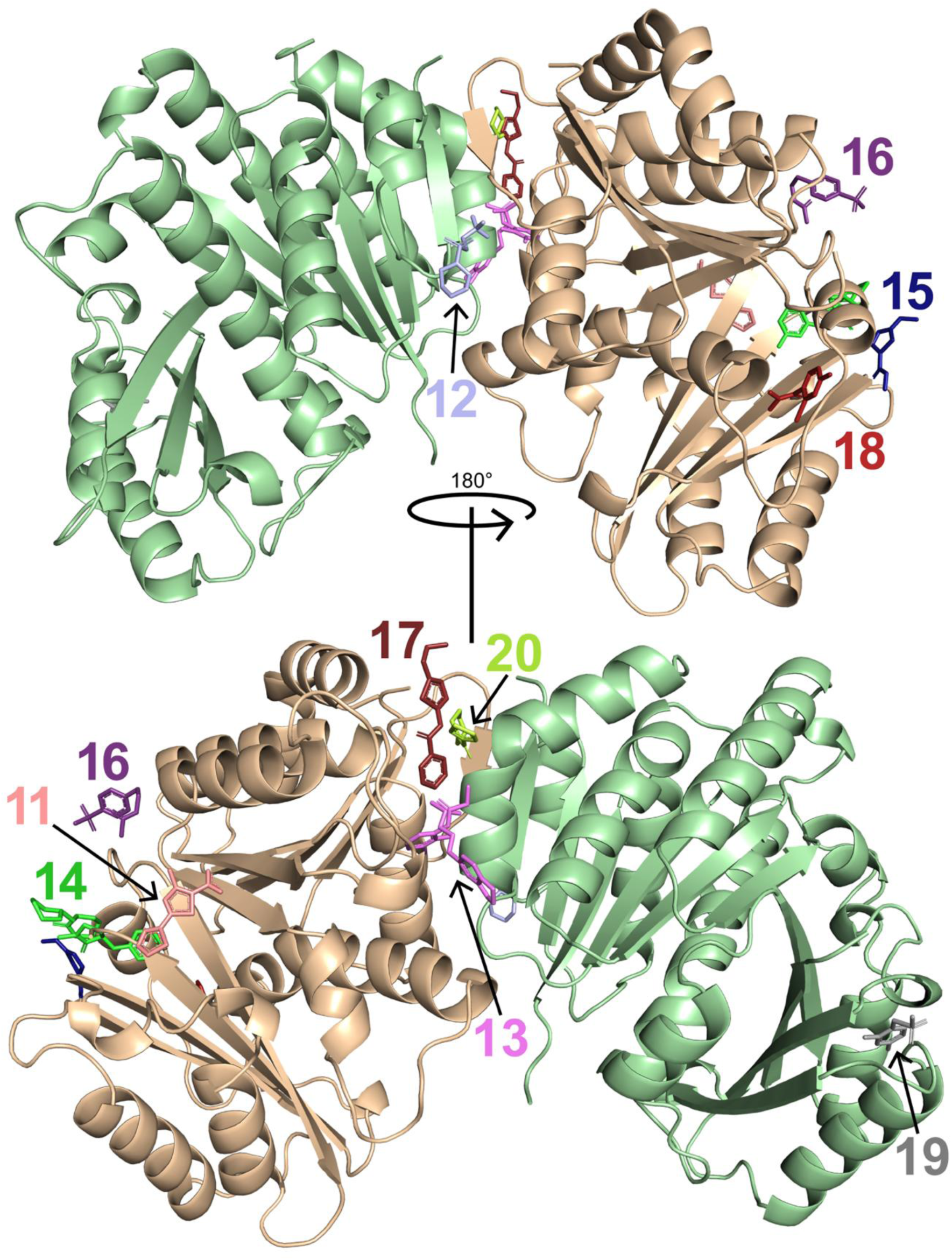
Locations of fragment-binding sites 11-20. Two views of the MtbFtsZ ground-state asymmetric unit are shown, related by a 180° rotation. Chain A (ON protomer) is pale green and chain B (OFF protomer) is wheat. Fragments are shown as sticks and colored according to sites ID.

**Figure S4.**
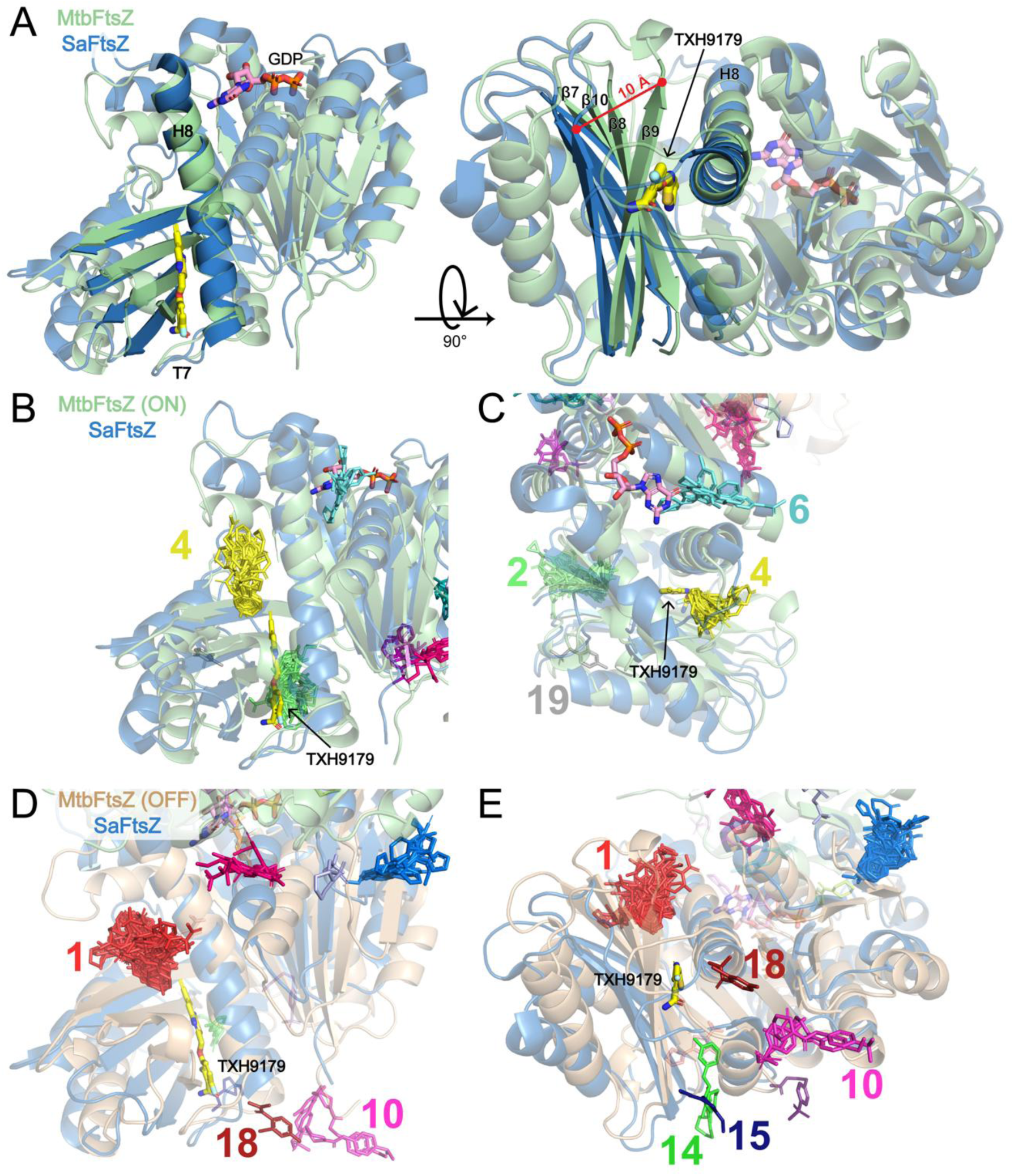
SaFtsZ (blue) allosteric binding pocket. **A**) Crystal structure of SaFtsZ complexed with TXH9179 (yellow) and GDP (pink) superposed to the ground state FtsZ model chain A (ON protomer; pale green). The structures are aligned by the core helix (H8 in MtbFtsz). **B-C**) SaFtsZ superposed onto MtbFtsZ chain A with hit fragments shown in sticks. **D-E**) SaFtsZ superposed onto the MtbFtsZ chain B (OFF protomer; wheat) with hit fragments shown in sticks.

---

**Supplementary Table S1. PDB accession codes and crystallographic information for MtbFtsZ fragment-screening structures.**

ftsz_supplemental_table1_20260824.csv

Each fragment-bound structure obtained from the crystallographic fragment is listed, the unmodelled datasets (pdb_000015aa) are listed last. For each structure, the screened fragment identifier, PDB chemical component ID, fragment-binding site(s), ligand chain and corresponding MtbFtsZ conformational state, refinement and crystallographic statistics are provided. Structures containing a fragment at more than one binding site list all observed sites and corresponding chain/state assignments. Enantiomers of fragments within the same dataset are separated into individual rows. All structures were deposited as part of PDB group deposition G_1002379.

**Supplemental File 1. DNA sequence of pET-28b(+)_MtbFtsZ**

pET-28b-plus_MtbFtsZ.txt

